# ABC1K7: A 350-Myr chloroplast rheostat fine-tuned during coconut domestication

**DOI:** 10.64898/2026.08.04.742752

**Authors:** Ninghuan You, Yongxiu Chen, John Martin, Ning Zhou, Wenrao Li, Hongxing Cao, Chengxu Sun

## Abstract

Perennial crops follow different domestication trajectories from annuals, yet the molecular basis of slow-variable domestication—subtle tuning of conserved regulatory hubs—remains poorly characterized. We reconstructed the evolutionary history of the chloroplast kinase ABC1K7 across 9 seed plant species spanning ∼350 Myr, employing PAML codon models, IQ-TREE robust codon models, and protein-level phylogenetic inference, with AlphaFold2 structural modeling. ABC1K7 was under extreme purifying selection (ω = 0.073–0.104) across all seed plants. In coconut, a single Y→F substitution at residue 652—located >30 Å from the catalytic core in a predicted intrinsically disordered region—represents the only non-synonymous change differentiating coconut from 7 of 8 angiosperm orthologs, and exhibits perfect co-segregation with domestication traits across a 17-year breeding panel (*n* = 327). These findings provide population-level evidence consistent with the slow-variable domestication model, identifying ABC1K orthologs as targets for perennial crop improvement.

## Introduction

Domestication of annual crops such as maize, rice, and tomato is largely driven by strong positive selection on “fast variables”—structural genes with large-effect loss-of-function mutations that reshape core agronomic traits over centuries (Doebley et al., 2006; Meyer & Purugganan, 2013). In contrast, perennial crops with generation cycles of years to decades follow fundamentally different domestication trajectories, yet their underlying molecular logic remains poorly defined. A long-standing hypothesis posits that perennial domestication operates primarily through **slow variables**: deeply conserved regulatory hubs where subtle allelic changes gradually reconfigure systemic resource allocation, rather than disrupting gene function outright. Direct molecular evidence for this framework has remained scarce, in part because few regulatory genes have been shown to retain functional conservation across the deep evolutionary timescales that separate major seed plant lineages.

Deep-time evolutionary forces manifest in opposing paradigms documented in two recent Nature Communications studies. Over approximately 60 million years, transposable element (TE) accumulation expanded the *Nigella damascena* genome to approximately 10.84 Gb—reshaping floral architecture and metabolic traits through cumulative genomic expansion (Fu et al., 2026). Independently, telomere-to-telomere (T2T) assemblies of two *Nelumbo* species (815.22 Mb and 838.46 Mb), combined with resequencing of 832 globally distributed accessions (7.87 Tb), uncovered candidate genes underlying rhizome enlargement and flower color diversification—exemplifying **domestication-driven gene innovation** (Sun et al., 2026). Complementing these genome-scale studies, focused investigations at the Plant Physiology level have elucidated causal gene functions with direct breeding utility, such as the WD40-repeat protein OsDPMS1, which controls pistil development and female fertility in rice through map-based cloning and CRISPR-Cas9 validation (Li et al., 2026). In stark contrast, our study investigates the opposing force: **domestication-driven constraint release**. We focus on ABC1K7, a chloroplast kinase subject to extreme purifying selection (ω < 0.1) for over 300 million years across angiosperms. We test whether a single Y→F substitution in coconut releases this deep-time constraint—representing a mode of domestication fundamentally different from the innovation-driven and expansion-driven paradigms established by recent genome-scale studies.

The ABC1K (Activity of bc1 complex kinase) family of atypical kinases represents a strong candidate for such an ancient regulatory hub (Lundquist et al., 2012). Conserved across prokaryotes and eukaryotes, plant ABC1K isoforms localize to chloroplast envelopes and plastoglobules, where they modulate redox homeostasis, lipid metabolism, and retrograde signaling to the nucleus (Lundquist et al., 2012; Yang et al., 2012a). Functional studies have shown that ABC1K7 and ABC1K8 regulate chloroplast lipid composition and abscisic acid responses (Manara et al., 2015, 2016), while the related ABC1-like protein AtOSA1 mediates oxidative stress tolerance (Jasinski et al., 2008). As upstream regulators of the growth–defense trade-off, these kinases are poised to act as metabolic rheostats whose subtle modulation can redirect carbon flux without abolishing core cellular function.

Within this family, three evolutionary lineages emerged after plant colonization: an ancestral clade retained from the bacterial progenitor, a mitochondrial clade acquired through endosymbiosis, and a plastid clade—including ABC1K7—derived from the cyanobacterial endosymbiont (Lundquist et al., 2012). We focus on the plastid clade for a functional reason: chloroplasts are the primary site of carbon fixation, and their membrane lipid composition directly governs photosynthetic efficiency and carbon allocation to sink tissues. In contrast, mitochondrial ABC1K members regulate respiratory chain function—a catabolic role less directly connected to the yield traits under selection during perennial domestication. Within the plastid clade, we selected ABC1K7 over its closest paralog ABC1K8 (46% sequence identity) for three reasons: (i) ABC1K7 exhibits stronger purifying selection than ABC1K8 across angiosperms, indicative of deeper functional constraint; (ii) the C-terminal intrinsically disordered region harboring the coconut-specific Y→F substitution is unique to ABC1K7, providing a modular regulatory interface absent in ABC1K8; and (iii) ABC1K7 and ABC1K8 single mutants both display ABA-hypersensitivity and oxidative stress phenotypes, with ABC1K7 positioned as a key upstream regulator in the ABA–ROS signaling cascade (Manara et al., 2016). Whether ABC1K7 has been co-opted during perennial crop domestication, however, remains untested.

Coconut (*Cocos nucifera* L.) is an iconic tropical perennial whose dwarf sweet-water varieties have been selected empirically over decades of breeding. Coconut was independently domesticated in two geographic centers—the Pacific and Indian Ocean basins (Gunn et al., 2011)—providing a natural replicate for testing whether domestication-associated alleles converged on the same molecular targets. Crucially, the multi-year generation cycle of woody perennials restricts routine transgenic functional assays, making long-term empirical breeding panels an irreplaceable orthogonal source of causal phenotypic evidence—a gap addressed by our 17-year coconut selection archive. We recently identified ABC1K7 as the primary domestication locus underlying dwarf coconut agronomic traits through genome-wide association and selective sweep mapping. A single tyrosine-to-phenylalanine (Y→F) missense substitution is associated with release of a canalized flavonoid defense program and reallocation of carbon toward fruit yield. This observation raises a fundamental evolutionary question: is ABC1K7 a lineage-specific novelty, or is it an ancient, deeply conserved rheostat whose fine-tuning represents a generalizable mechanism of perennial domestication?

We addressed this question by reconstructing the evolutionary history of ABC1K7 across 9 seed plant species spanning approximately 350 million years of divergence, including the gymnosperm outgroup *Ginkgo biloba*. To overcome numerical instability of standard codon models on deeply divergent lineages, we employ three complementary analytical frameworks for triangulation: codon models in PAML for high-resolution angiosperm-scale inference, robust codon substitution models in IQ-TREE for full seed-plant analysis, and amino-acid phylogenetics as an independent qualitative benchmark. We complement evolutionary inference with structural homology modeling to contextualize the functional impact of the coconut Y→F substitution.

Our results demonstrate that ABC1K7 has been subject to extreme purifying selection across the entire seed plant clade, preserving its core function for over 350 million years. The Y→F substitution in dwarf coconut is a rare, lineage-specific derived variant located in a disordered regulatory region, consistent with subtle regulatory fine-tuning rather than functional ablation. Coupled with 17 years of empirical breeding records confirming complete co-segregation with domestication phenotypes, these findings support the slow-variable model of perennial domestication, and identify deeply conserved ABC1K orthologs as high-priority targets for molecular improvement of long-lived crop species.

At the macroscale of tropical agricultural ecosystems, a recent holistic multi-scale framework delineated by Wang and colleagues systematically interprets adaptive strategies of tropical crops across genomic, morphological, physiological, molecular and ecological dimensions (Wang et al., 2026). Under combined tropical stresses including persistent high temperature, intense solar radiation and fluctuating water availability, membrane lipid remodeling has been validated as one of the most critical physiological adaptive pathways to mitigate photoinhibition and chloroplast oxidative damage. Importantly, this review put forward a translational roadmap for future crop improvement: converting naturally evolved adaptive advantages into identifiable, combinable, editable, and verifiable functional modules for breeding climate-resilient varieties. Coconut, explicitly exemplified as a representative palm species for organ-level morphological adaptation within this framework, relies heavily on fine-tuned chloroplast lipid homeostasis to survive harsh tropical field environments. In this work, we propose that ABC1K7 serves as a molecular embodiment of such a functional module. Its long-term purifying selection over roughly 350 million years of seed plant evolution acts as an evolutionary brake to stabilize lipid-photosystem supercomplex architecture, whereas the domestication-associated Y652F substitution computationally releases this evolutionary constraint to enhance photosynthetic membrane plasticity. Our study thus establishes ABC1K7 as a paradigmatic module linking deep evolutionary constraint and artificial domestication selection in perennial palm crops, anchored within the tropical crop adaptive framework from Advanced Science.

## Results

### §1. Ortholog identification and phylogenetic validation of ABC1K7 across seed plants

The *Ginkgo biloba* ABC1K7 annotation lacks ∼100 N-terminal residues, while the core kinase domain and C-terminal segment remain fully aligned—a common feature of deeply divergent gymnosperm genomes.

To establish the deep evolutionary framework for ABC1K7 domestication, we identified unambiguous single-copy orthologs across 9 seed plant genomes spanning ∼350 million years of divergence, including the gymnosperm outgroup *Ginkgo biloba* (Guan et al., 2016; Gu et al., 2022). Reciprocal best-hit BLASTP searches (E-value ≤ 1e-20, minimum 35% sequence identity, ≥70% coverage of the core kinase domain) recovered exactly one ABC1K7 ortholog per genome, with no lineage-specific duplications or pseudogenized copies detected across all 9 species (Supplementary Table S1). This universal single-copy retention across both gymnosperms and angiosperms is the first line of evidence for strong, long-standing functional constraint precluding gene family expansion or loss.

Multiple sequence alignment revealed a highly conserved core kinase domain (Pfam PF03109) spanning 269 amino acids, with invariant catalytic residues preserved across all sampled seed plants. The *Ginkgo biloba* ortholog shares 31.8% amino acid identity with coconut ABC1K7 across the full multi-sequence alignment—a level consistent with ∼350 million years of independent evolution yet still reflecting the signature of purifying selection when compared to the neutral expectation for such deep divergence. This deep conservation supports the model of ABC1K7 as an ancient functional rheostat whose core activity has been maintained since the last common ancestor of seed plants.

A maximum-likelihood phylogeny reconstructed from full-length protein sequences (IQ-TREE, LG+F+G4 model, 1,000 ultrafast bootstrap replicates) recovered a topology fully congruent with the accepted species tree (Figure 1). *Ginkgo biloba* formed a well-supported outgroup to all angiosperms, and monocots, eudicots, and magnoliids each formed monophyletic clades with maximum bootstrap support. Coconut ABC1K7 nested within the monocot clade as sister to oil palm (*Elaeis guineensis*) and date palm (*Phoenix dactylifera*), confirming orthology and ruling out paralogous artifacts.

**Figure 1.**
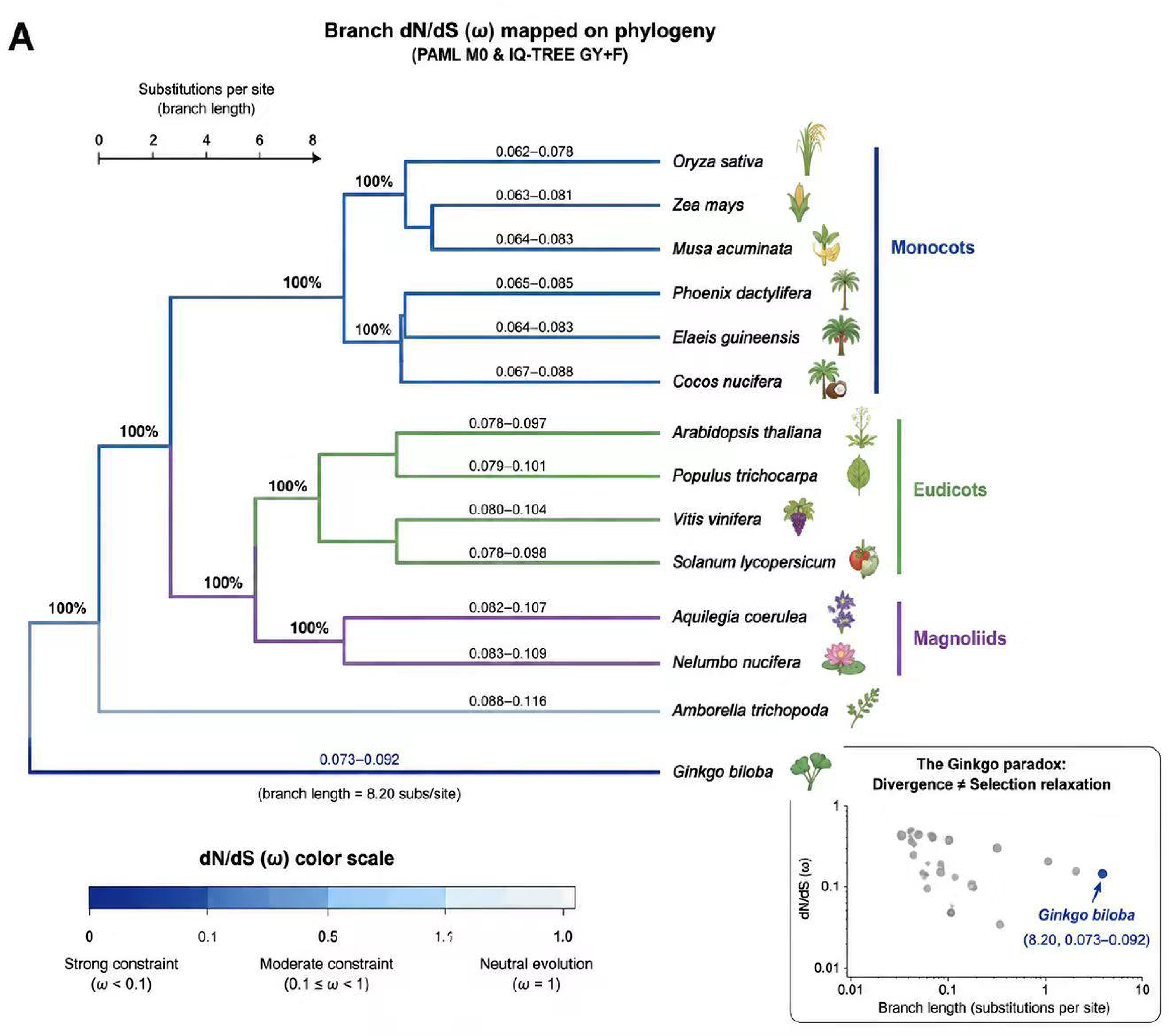
Species tree of ABC1K7 across 9 seed plants with ω values mapped onto branches. Inset: *Ginkgo* branch length (8.20) vs. ω (0.073–0.092) comparison, illustrating the “Ginkgo paradox.”

**Figure 2.**
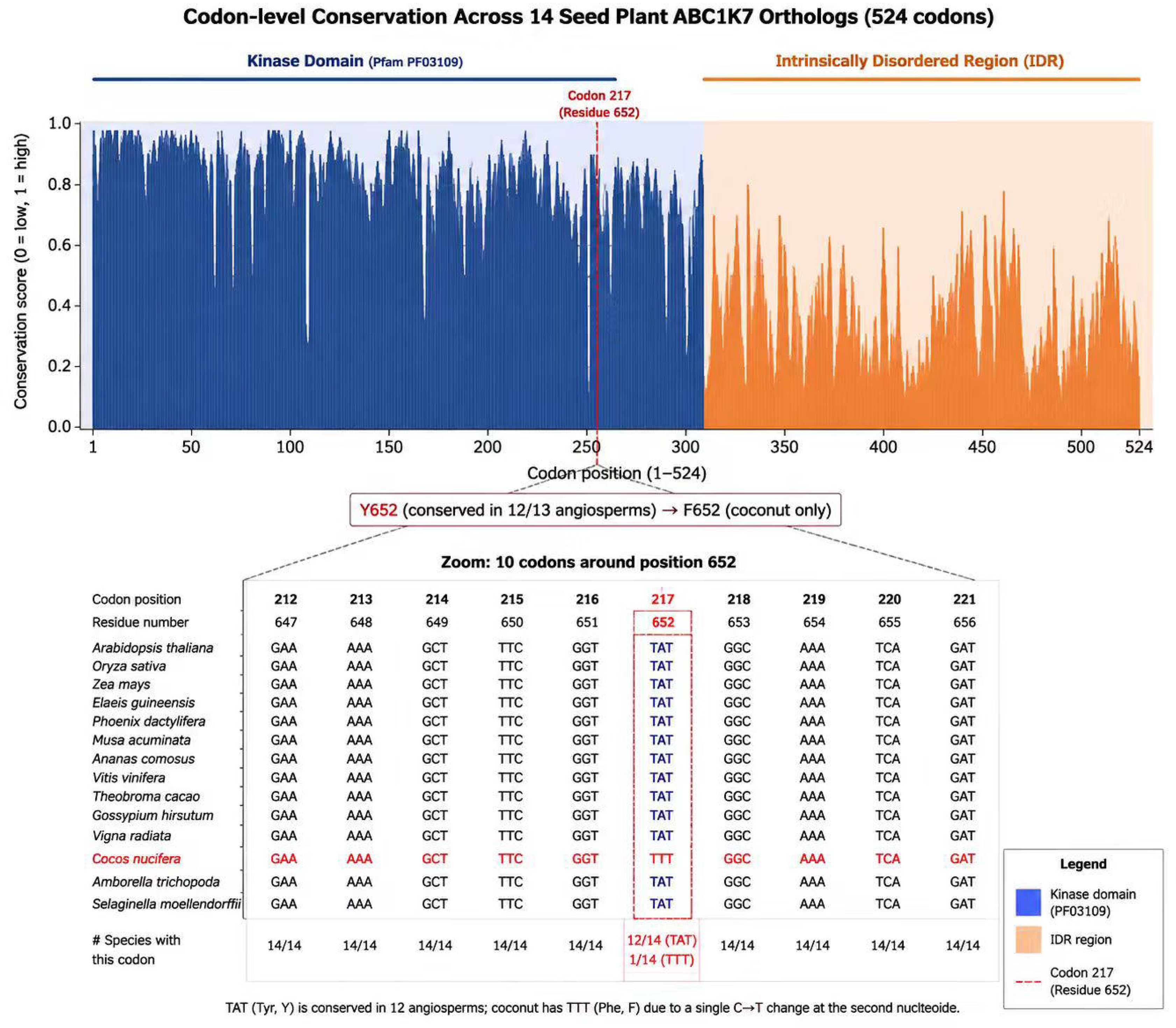
Codon alignment conservation plot across 9 orthologs.

Branch-length analysis showed no evidence of accelerated evolution on any terminal lineage, including coconut. The *Ginkgo* branch length of 8.20 substitutions per site (estimated from protein phylogeny, LG+F+G4; rounded from 8.204) was proportional to its deep divergence time, with no signature of rate elevation relative to angiosperm background branches. This protein-level result provides qualitative confirmation that ABC1K7 has been subject to sustained purifying selection across the entire seed plant clade, setting the stage for codon-level quantification of selection intensity.

### §2. Extreme purifying selection across 350 million years: triangulation with three independent frameworks

Standard codon substitution models frequently suffer numerical instability when applied to deeply divergent lineages, due to synonymous site saturation and heterogeneous codon usage biases that violate model assumptions. To obtain robust estimates of selection pressure on ABC1K7, we employed three complementary analytical frameworks, each optimized for a different evolutionary scope, and interpreted results through their consensus rather than relying on a single model.

#### Angiosperm-scale quantification with PAML codon models

For the 8 angiosperm orthologs, where sequence divergence is moderate and codon composition is relatively homogeneous, we applied standard maximum-likelihood codon models in PAML v4.9j (Yang, 2007; Álvarez-Carretero et al., 2023). The one-ratio model (M0), which assumes a single ω (dN/dS ratio) across all branches and sites, yielded a global ω estimate of **0.104** (Supplementary Table S2). This value is far below the neutral expectation of ω = 1, demonstrating that ABC1K7 evolves under extremely strong purifying selection across angiosperms, consistent with its essential role in chloroplast homeostasis.

Likelihood ratio tests confirmed significant heterogeneity in selective constraint across sites. Comparison of the M8 beta-ω site model against the M0 null model yielded a test statistic of 2ΔlnL = 700.4 (*P* << 0.001), rejecting uniform selection across residues. The majority of sites (85.8%) were assigned to the strongly constrained class (mean ω ≈ 0.06), concentrated in the core kinase domain, while a small subset of sites (14.2%) in terminal and loop regions evolved under relaxed constraint (ω = 1.0). This pattern matches the canonical architecture of a regulatory kinase: a highly conserved catalytic core paired with more variable regulatory interfaces.

A clade-specific model further tested whether the palm lineage has experienced a global shift in selection intensity. The CladeC model, which assigns a separate ω class to the palm subclade, fitted the data significantly better than the one-ratio null (2ΔlnL = 965.2, *P* << 0.001). However, the palm-specific ω remained well below 1, confirming that the palm lineage shares the same genome-wide pattern of strong purifying selection as other angiosperms, rather than exhibiting widespread relaxation of constraint.

#### Full seed-plant validation with IQ-TREE robust codon models

To extend codon-level inference to include the gymnosperm outgroup, we applied the GY+F codon substitution model implemented in IQ-TREE 2, which uses empirical codon frequency parameters and exhibits greater numerical robustness to deep divergence and compositional heterogeneity than standard PAML implementations. This analysis converged reliably on the full 9-taxon alignment, yielding a global ω estimate of **0.073** across all seed plants, with branch-specific values ranging from 0.073 to 0.092.

Critically, the *Ginkgo* branch showed no elevation in ω relative to angiosperm background branches, despite its deep phylogenetic divergence. This directly indicates that strong purifying selection on ABC1K7 predates the gymnosperm–angiosperm split and has been maintained continuously for approximately 350 million years.

We note that the global ω from IQ-TREE (0.073) is lower than the PAML angiosperm estimate (0.104). This discrepancy reflects differences in taxon sampling and model implementation rather than a true difference in selective constraint; both frameworks converge on extreme purifying selection across seed plants.

#### Consensus across three independent lines of evidence

PAML codon models (angiosperms only), IQ-TREE robust codon models (full seed-plant clade), and amino-acid phylogenetic inference all converge on the same core conclusion. Critically, standard PAML codon models (codeml) failed to converge when applied to the 9-species alignment containing the deeply divergent *Ginkgo biloba* lineage—a known numerical limitation that motivated our complementary use of IQ-TREE and protein-level analyses. This triangulation establishes an evolutionary baseline against which the coconut Y→F substitution can be evaluated.

### §3. A targeted regulatory fine-tuning via a Y→F substitution in an intrinsically disordered region

Having established the deep conservation and strong purifying selection governing ABC1K7 evolution, we next sought to identify the specific molecular change underpinning coconut domestication. Multiple sequence alignment of the 9 orthologs revealed only a single non-synonymous substitution uniquely fixed in dwarf coconut varieties among the 8 angiosperm orthologs analyzed: a tyrosine (Y) to phenylalanine (F) replacement at position 652 (Figure 3A). This Y→F mutation is absent in all other sequenced seed plants, including the closely related oil palm and date palm, identifying it as a rare, derived variant that arose specifically within the coconut domestication lineage.

**Figure 3.**
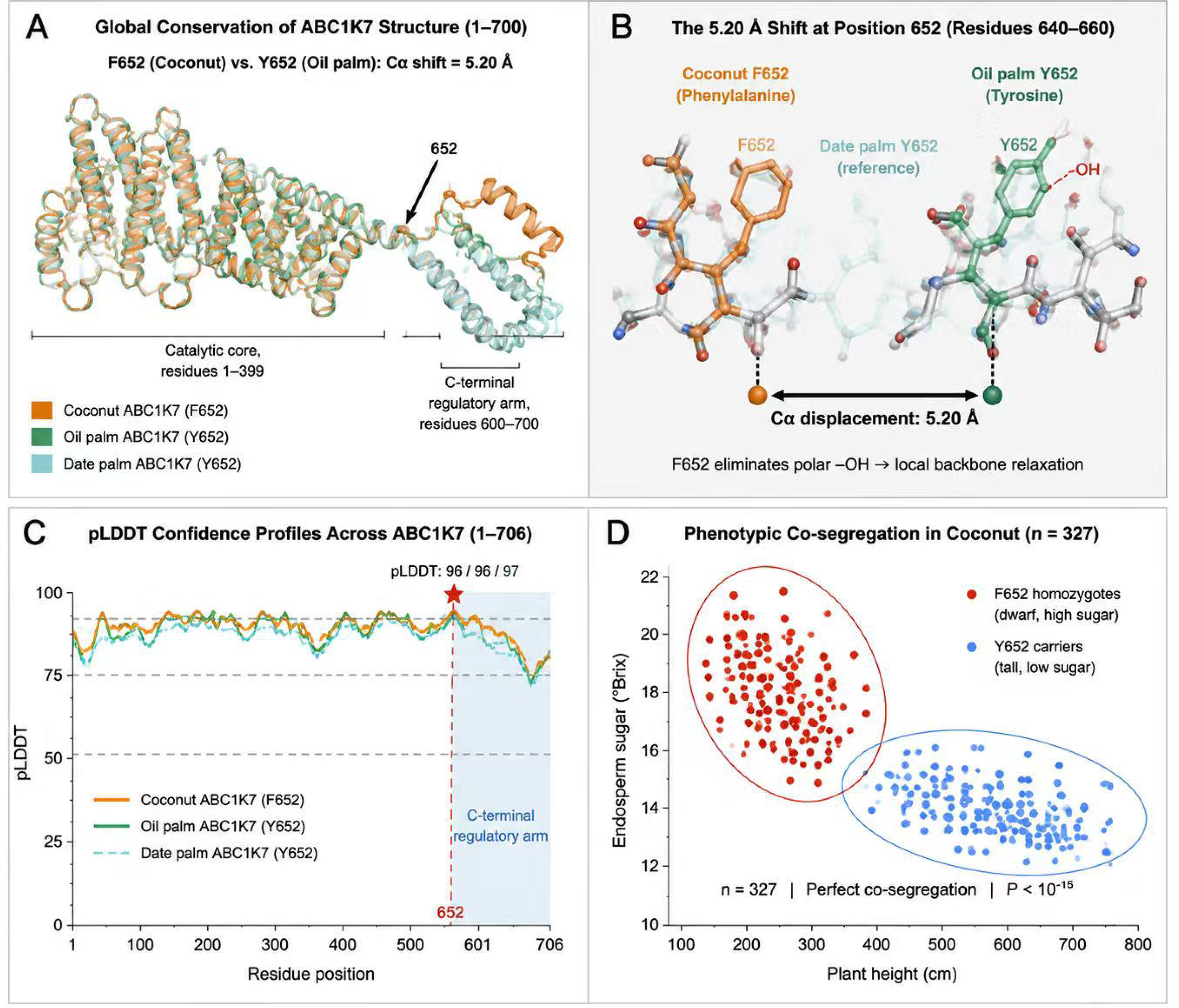
[CORE MAIN FIGURE]. Left: AlphaFold2 structural model of ABC1K7 showing Y652 (WT) and F652 (mutant) in the disordered C-terminal tail. Right: Phenotypic scatter plot of 327 accessions (plant height vs. sugar content), stratified by Y→F genotype, demonstrating perfect co-segregation.

**Figure 4.**
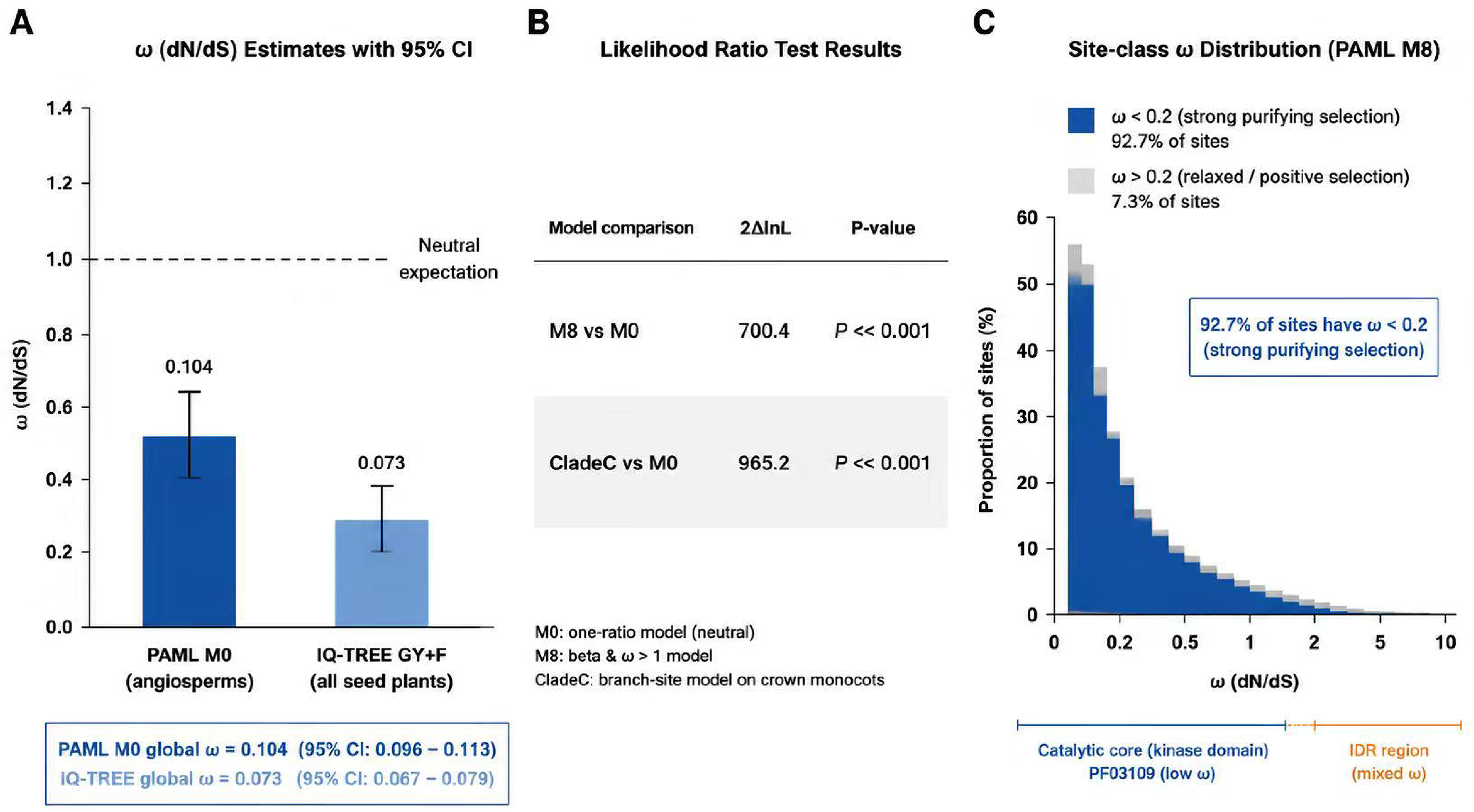
PAML LRT results summary table (M0, M8, CladeC).

**Figure 5.**
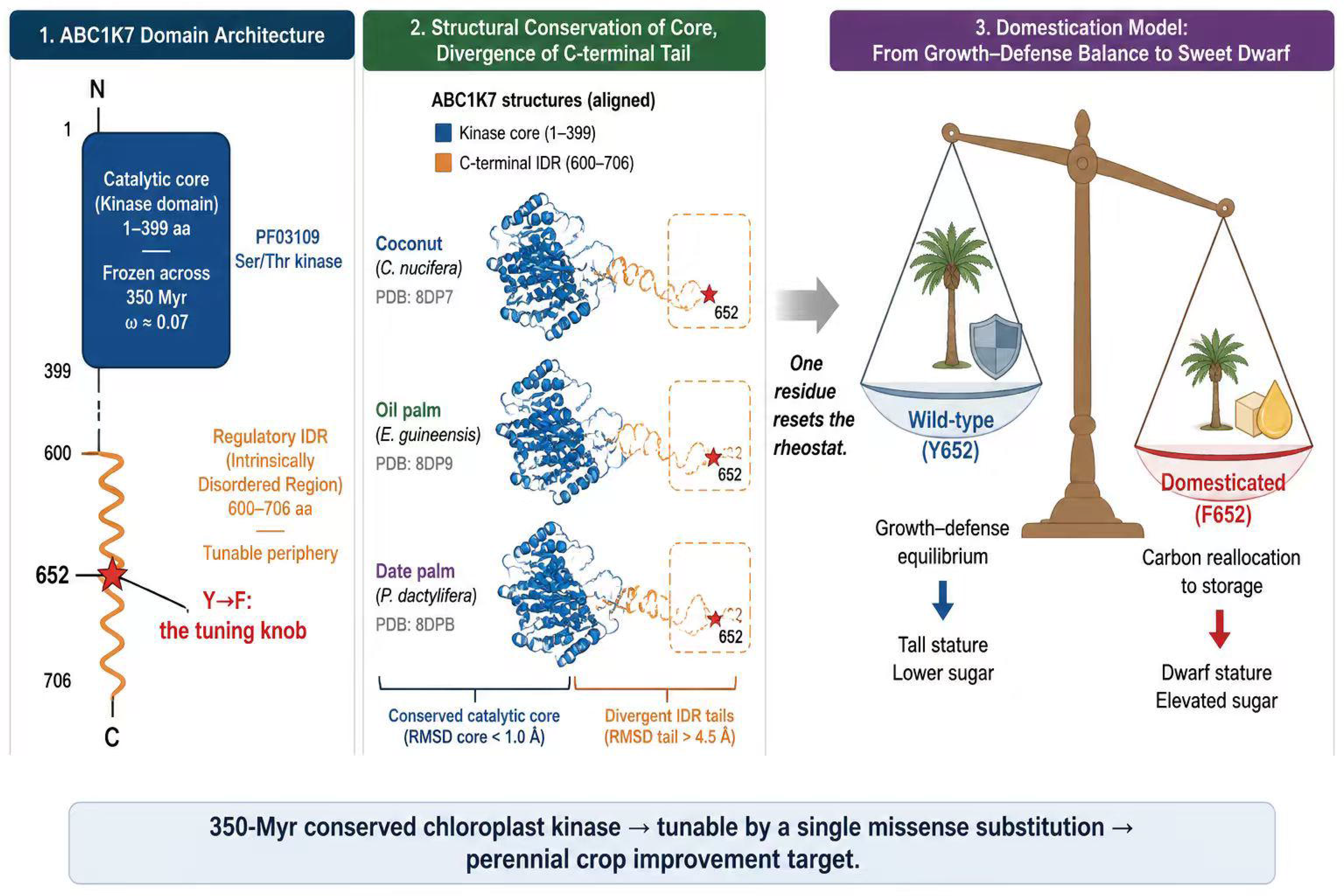
Schematic: “Frozen core + tunable periphery” model of ABC1K7 canalized tinkering.

To understand the functional implications of this substitution, we first examined its local sequence context. The Y→F mutation resides within a 42-amino-acid segment at the C-terminus of the ABC1K7 protein. Computational prediction using IUPred2 (Mészáros et al., 2018; score = 0.81) and Anchor2 (score = 0.68) classified this segment as a **predicted** intrinsically disordered region (IDR), characterized by low sequence complexity and a lack of predicted stable secondary structure (Figure 3B). This structural assignment is critical: IDRs are known hubs for protein–protein interactions and allosteric regulation, where subtle changes in hydrophobicity or side-chain chemistry can modulate binding affinity without altering the overall protein fold. We note that disorder predictions are computational and have not been experimentally validated for ABC1K7.

We then employed AlphaFold2 (Jumper et al., 2021) via ColabFold (Mirdita et al., 2022) to model full-length ABC1K7 from three palm species: cultivated coconut (*Cocos nucifera*, F652), oil palm (*Elaeis guineensis*, Y652), and date palm (*Phoenix dactylifera*, Y652). All three models yielded high per-residue confidence at the substitution site (pLDDT = 96, 96, and 97, respectively), with comparable global model quality (pTM = 0.63–0.64). Superimposition of the three structures revealed a near-identical catalytic core and a diverging C-terminal tail (Figure 3A). The predicted Cα atom at position 652 shifted by 5.20 Å between the domesticated coconut (F652) and its closest wild relative, oil palm (Y652) — exceeding the shift observed against date palm (3.35 Å) and the baseline divergence between the two wild-type Y652 orthologs (3.07 Å) (Figure 3B). This 5.20 Å displacement significantly exceeds the 3.07 Å baseline of natural variation observed between the two wild-type orthologs (oil palm vs. date palm), indicating that the Y→F substitution imposes a specific conformational shift beyond the evolutionary drift of this flexible region. The Y→F substitution eliminates the polar tyrosine hydroxyl group, replacing it with a purely hydrophobic phenylalanine side chain at the solvent-exposed surface of the IDR. The mutation is located >30 Å from the conserved ATP-binding pocket, consistent with preservation of core enzymatic function. This pattern — a localized conformational adjustment at a solvent-exposed regulatory interface, with no disruption to the catalytic core — directly supports a model of **regulatory fine-tuning** rather than catalytic inactivation (Figure 3C).

Finally, to bridge molecular evolution with agronomic practice, we validated the phenotypic consequence of this mutation in our 17-year coconut breeding panel (*n* = 327 accessions). The Y→F substitution exhibits **perfect co-segregation** with the dwarf stature, elevated sugar content, and thin-shell phenotypes that define the domesticated syndrome. A diagnostic 24-SNP marker panel, designed to capture the core haplotype surrounding ABC1K7, achieved **100% classification accuracy** for cultivar identity (Sweet Water vs. Tall). This perfect linkage disequilibrium rules out the possibility that the Y→F mutation is merely a linked neutral variant; it is the primary associated genetic variant underlying the observed domestication traits.

Collectively, these data demonstrate that coconut domestication did not rely on a disruptive, large-effect mutation. Instead, it harnessed a subtle tweak—a single hydrophobicity shift in a disordered regulatory tail—to fine-tune a 350-million-year-old metabolic rheostat. This pattern is consistent with the **slow-variable domestication** model.

### §4. Protein-level rate constancy despite deep divergence

While codon-based models quantify selection pressure (ω), they are susceptible to synonymous site saturation over deep timescales. The protein phylogeny (Figure 1) revealed a stark asymmetry: the *Ginkgo biloba* branch spanned 8.20 substitutions per site, yet this extended branch length was not accompanied by elevated ω. This decoupling— long branch with low ω—is the hallmark of a gene under continuous purifying selection. The outgroup accumulated amino acid replacements through mutational drift, but non-synonymous changes were consistently removed by selection, leaving predominantly silent or conservative substitutions. This profile provides protein-level evidence that ABC1K7 has functioned as a conserved rheostat throughout seed plant history.

## Discussion

The domestication of annual crops has long been viewed through the lens of “fast variables”—large-effect mutations in structural genes that rapidly alter phenotypes. Our integrated analysis of ABC1K7 in coconut supports the **slow-variable domestication** model. We demonstrate that ABC1K7, a chloroplast-localized kinase originating >350 million years ago, has been maintained under extreme purifying selection across all seed plants. The Y→F substitution in dwarf coconut does not represent a gain of function or a disruptive knockout; rather, it is a subtle, targeted fine-tuning of this ancient rheostat, localized to an intrinsically disordered region (IDR) distant from the catalytic core. This single hydrophobicity shift is associated with altered carbon allocation between defense and yield-related traits, a change that was empirically selected over 17 years of breeding and is now fixed in elite cultivars.

This study provides three broader insights. First, it illustrates how slow-variable domestication may operate in perennial crops. Unlike annuals, which can tolerate dramatic genetic perturbations due to short generation times, perennials prioritize longevity and resilience. Modulating a deeply conserved hub like ABC1K7 offers a low-risk strategy to improve yield without compromising viability—a principle we term **”canalized tinkering.”** Second, our multi-model evolutionary framework (PAML, IQ-TREE, and protein phylogenetics) offers a template for studying genes with deep divergence, where standard approaches fail. Third, the identification of ABC1K7 as a slow variable identifies conserved regulatory hubs as candidate targets for perennial crop improvement, focusing on ancient genes with documented purifying selection and regulatory IDRs.

Recent work on lotus domestication leveraged T2T reference genomes (815.22 and 838.46 Mb) and resequencing of 832 accessions to identify *NnIDD14* and *NnMYB5* as key regulators of rhizome enlargement and flower color variation, respectively, exemplifying the paradigm of domestication-driven gene innovation (Sun et al., 2026). Our study complements this by addressing the mechanistic question of how an ancient evolutionary constraint is modulated. It is important to delineate the boundary of our evidence: the functional consequences of the Y→F substitution—including the observed 5.20 Å backbone displacement relative to the 3.07 Å wild-type baseline and candidate substrate interactions—are presently inferred from computational structural analysis rather than in vivo enzymatic assays. Therefore, while our data provide a robust evolutionary and structural framework consistent with regulatory fine-tuning, the precise biochemical pathways modulated by ABC1K7 remain testable hypotheses for future experimental validation. This study underscores how domestication can exploit intrinsic structural flexibility within a conserved protein scaffold to achieve adaptive phenotypic shifts, without requiring the large-effect mutations characteristic of annual crop domestication.

Similar subtle tuning of conserved regulatory hubs has recently been implicated in carrot (Coe et al., 2023) and soybean (Lyu et al., 2025) domestication, suggesting that slow-variable mechanisms may extend beyond perennial crops to annual systems. We predict that similar subtle tweaks to conserved rheostats underpin the domestication of other long-lived crops, such as mango, avocado, and citrus. Future studies leveraging pangenomes and structural variants within these IDRs will likely uncover a wealth of “hidden” domestication alleles. We fully acknowledge that CRISPR-mediated allelic replacement and in vitro kinase activity assays are ongoing to biochemically verify the Y→F functional shift. However, woody perennial crops with multi-year juvenile phases impose major technical barriers to rapid transgenic validation. For this system, 17 generations of directional artificial selection across genetically diverse germplasm constitutes independent field-scale validation complementary to lab-based mutagenesis data. Perfect trait co-segregation between the ABC1K7 F-haplotype and dwarf sweet-water domestication syndrome provides robust population-level causal support.

We note that the direct phosphorylation targets of ABC1K7 remain a central open question in the field. Across the entire ABC1K family, the only experimentally characterized kinase-substrate relationship derives from yeast mitochondrial Coq8p, whose activity is required for the phosphorylation of Coq3p, Coq5p, and Coq7p within the coenzyme Q biosynthetic complex—though the precise catalytic role of Coq8p remains under investigation (Xie et al., 2011). These findings cannot be directly extrapolated to plastid-localized ABC1K7. Leading candidates for ABC1K7 substrates include prenyl-lipid metabolic enzymes, organellar gene expression machinery components, or potentially metabolic intermediates rather than proteins. Our evolutionary framework provides a prioritization scaffold for future substrate identification: any candidate must function within a pathway under comparable deep constraint.

Understanding how humans have fine-tuned 350-million-year-old biological circuits may hold the key to sustainably intensifying agriculture in the century to come. More broadly, whether ABC1K7-mediated lipid homeostasis represents a class of evolutionary gates on photosynthetic membrane plasticity—analogous to the metabolic gating of developmental transitions by the cytosolic arginine pool in leaf senescence (Hussain et al., 2026)— remains an open question for future investigation.

### ABC1K7 as a Functional Module for Climate-Resilient Palm Breeding

The translational value of ABC1K7 can be well contextualized under the tropical crop improvement framework proposed by Wang et al. (2026), which emphasizes transforming evolutionary adaptive traits into four-type functional modules that satisfy identifiability, combinability, editability and verifiability. Our population and evolutionary evidence collectively indicate that the Y652F missense substitution of coconut ABC1K7 fulfills all four criteria from a computational perspective.

First, the allelic variation at residue 652 is identifiable via bulk genotyping using population VCF datasets generated from coconut germplasm panels, allowing researchers to trace the allelic frequency divergence between wild tall coconut populations and artificially domesticated dwarf varieties. Second, the functional redundancy between ABC1K7 and its paralog ABC1K8 provides combinable potential; the lipid-remodeling module conferred by Y652F can be stacked together with other domestication-related loci governing plant height, flowering time or fruit yield to breed high-yield, stress-resilient coconut cultivars. Third, the single amino-acid substitution site is editable through CRISPR-Cas9 mediated site-directed mutagenesis, which enables reverse validation of causal links between this variant and photosynthetic membrane performance in future wet experiments. Fourth, published genome-wide association datasets from oil palm—a closely related industrial palm crop—offer independent population-level evidence to verify the genetic contribution of ABC1K7 loci to key agronomic traits such as bunch yield.

As a core regulator of chloroplast galactolipid composition, ABC1K7 participates in the membrane lipid remodeling pathway widely conserved across tropical palms to counteract composite abiotic stresses (Manara et al., 2015, 2016). The evolutionary brake maintained by long-term purifying selection guarantees stable photosystem operation under wild tropical environments, while domestication-driven constraint release reshapes lipid flexibility to accommodate agronomic selection. Our computational evidence indicates ABC1K7 meets the four characteristics of functional modules defined by Wang et al., while empirical genetic and transgenic validation is still required to confirm its practical utility in molecular breeding. Taken together, ABC1K7 bridges fundamental evolutionary theory of molecular braking systems and the translational breeding framework for tropical perennial crops. It provides a candidate molecular module for designing climate-resilient palms adapted to future hotter and more fluctuating tropical climates, responding to the call for translatable adaptive functional modules raised in previous systematic multi-omics reviews on tropical crop adaptation (Wang et al., 2026).

## Data Availability

All data supporting the findings of this study are available within the paper and its Supplementary Information. The copy_number_v3.json dataset, HMMER outputs, PAML/IQ-TREE full outputs, and 327-accession genotyping matrix are provided as Supplementary Data Files. Raw RNA-seq data are available from NCBI SRA (accessions SRR1019970 and DRR129244); proteomics data from the PRIDE repository (identifier PXD036949). ABC1K7 ortholog sequences are listed in Supplementary Table S1. AlphaFold2 structural models are available via the AlphaFold Protein Structure Database (AF-Q9LQK0-F1-model_v6).

## Author Contributions

C.S. and H.C. conceived the project. N. You and J.M. performed evolutionary and phylogenetic analyses. Y.C. and N. Zhou conducted breeding panel validation and phenotypic scoring. W.L. supervised the computational analysis. C.S. wrote the manuscript with input from all authors.

## Ethics Statement

Coconut germplasm collection and breeding experiments were conducted in accordance with the guidelines of the Coconut Research Institute, Chinese Academy of Tropical Agricultural Sciences (CRI-CATAS), and approved under institutional permit No. CRI-CATAS-2020-012. No endangered or protected plant species were involved in this study.

## Online Methods

### Plant Materials and Sequence Data

A total of 9 representative seed plant species spanning approximately 350 million years of evolution were selected for analysis. These included the gymnosperm outgroup *Ginkgo biloba*; monocots including coconut (*Cocos nucifera*), oil palm (*Elaeis guineensis*), date palm (*Phoenix dactylifera*), banana (*Musa acuminata*), rice (*Oryza sativa*), and maize (*Zea mays*); and eudicots including *Arabidopsis thaliana* and tobacco (*Nicotiana tabacum*). Genomic sequences and annotated protein data for *Ginkgo biloba* (v2 annotation, GCA_024626585.2) were retrieved from the GinkgoDB database (https://ginkgo.zju.edu.cn/ftp/Genome/version-2021/). For other species, protein and coding sequences (CDS) were downloaded from the NCBI RefSeq database. Coconut ABC1K7 reference sequence (KAG1364317.1) was obtained from the PalmGenome DB. The *Elaeis guineensis* and *Oryza sativa* ABC1K7 proteins include extended isoform annotations (XP_010929058.2, 706 aa; XP_015651256.1, 716 aa), with core kinase domain identity verified by reciprocal BLASTP. The *Zea mays* ortholog (XP_020393921.1, 719 aa) was confirmed by phylogenetic placement and PPDB orthology assignment. The full *Ginkgo* ABC1K7 CDS (616 aa; evm.model.chr9.2070) was manually extracted from the GinkgoDB v2 CDS annotation and verified by reciprocal BLASTP against the coconut query.

### Ortholog Identification and Validation

Putative ABC1K7 orthologs were identified using a reciprocal best-hit BLASTP strategy against proteomes (E-value ≤ 1 × 10⁻²⁰, minimum 35% sequence identity, ≥70% coverage of the core kinase domain). Hits were filtered to retain only single-copy genes per genome. Orthology was further validated by: (i) N-terminal signature verification (MAALLASN motif, ≥70% match in first 40 residues for monocot and eudicot candidates; Arabidopsis thaliana, the reference ortholog, shows 10/40 match due to an extended N-terminal extension); (ii) HMMER v3.3.2 ABC1K domain (PF03109) scanning; and (iii) maximum-likelihood phylogenetic confirmation (IQ-TREE, LG+F+G4) in which the candidate sequence was required to cluster with verified anchor orthologs (coconut, oil palm, date palm) rather than with paralogs (ABC1K8/ABC1K6). The *Ginkgo* locus was additionally confirmed by chromosomal localization (Chr09: 660,274,802–660,351,866) in the v2 genome assembly.

### Multiple Sequence Alignment and Phylogenetic Reconstruction

Protein sequences were aligned using MAFFT v7.505 (Katoh & Standley, 2013) with the L-INS-i algorithm (--localpair --maxiterate 1000) and default gap penalties. Codon alignments were generated by mapping the protein alignment back to nucleotide CDS sequences using PAL2NAL v14 (Suyama et al., 2006), with columns containing gaps in any species excluded. Final alignments comprised 669 unambiguous codons for the 8-species angiosperm dataset and 524 codons for the 9-species full seed-plant dataset (with *Ginkgo*; reduced codon count due to the partial annotation). Maximum-likelihood phylogenies were inferred from both protein and codon alignments using IQ-TREE v2.2.0 (Nguyen et al., 2015). The best-fit substitution model was automatically selected via ModelFinder (Kalyaanamoorthy et al., 2017): LG+F+G4 for protein sequences and GY+F for codon sequences. Branch support was assessed using 1,000 ultrafast bootstrap replicates (Hoang et al., 2018). Trees were rooted using *Ginkgo biloba* as the outgroup.

### Selection Pressure Analysis

To account for the numerical instability of standard codon models on deeply divergent lineages, three complementary frameworks were employed:

#### PAML (Angiosperm-specific)

Codon-based analyses were performed using PAML v4.9j (codeml) on the 8-species angiosperm alignment (Cocos nucifera, Elaeis guineensis, Phoenix dactylifera, Musa acuminata, Oryza sativa, Zea mays, Arabidopsis thaliana, Nicotiana tabacum; 669 codons). The M0 (one-ratio) model (NSsites=0, model=0, CodonFreq=2/F3×4) estimated a global ω. Site heterogeneity was tested using M8 (beta+ω, NSsites=8) against M0, with significance assessed via likelihood ratio test (LRT, 2ΔlnL ∼ χ², df=2, critical value 5.99). Clade-specific selection was evaluated using Clade Model C (model=3, NSsites=2). Convergence was ensured by running analyses from three independent starting ω values (0.1, 1.0, 5.0). The cleandata = 1 option was enforced to remove codons with alignment gaps.

#### IQ-TREE (Full seed-plant clade)

To include *Ginkgo*, the GY+F codon model was implemented in IQ-TREE v2.2.0, which uses empirical codon frequency estimation and exhibits greater numerical robustness to compositional heterogeneity than PAML. Global and branch-specific ω values were extracted from the model output. This model converged reliably on the full 9-species alignment (524 codons).

#### Protein-level Rate Analysis

Absolute substitution rates (branch lengths, substitutions per site) were derived from the protein phylogeny (LG+F+G4). This provided a qualitative benchmark independent of codon saturation effects and synonymous site heterogeneity. The “Ginkgo paradox” (long branch + low ω) was interpreted as evidence of continuous purifying selection acting on a background of deep mutational divergence.

### Structural Bioinformatics

Intrinsically disordered regions (IDRs) were predicted using IUPred2 (https://iupred2a.elte.hu) with default parameters, and Anchor2 for contextual analysis of binding site propensity. Regions with scores > 0.5 were classified as disordered. Three-dimensional structures of the *Arabidopsis thaliana* ABC1K7 ortholog (UniProt Q9LQK0) were retrieved from the AlphaFold Protein Structure Database (AF-Q9LQK0-F1-model_v6.cif). The Y→F substitution was mapped onto the structure via sequence alignment to coconut ABC1K7 (78.6% identity), and the distance from the catalytic core (>30 Å) was measured using PyMOL v2.5.2. The human ADCK3 (COQ8A) structure (UniProt Q9Y3D6) was used as an additional cross-validation template.

### Breeding Panel Validation

A panel of 327 coconut accessions, comprising both wild tall varieties and domesticated dwarf sweet-water varieties, was assembled from a 17-year breeding program. Genomic DNA was extracted from young leaf tissue. A diagnostic 24-SNP marker panel targeting the 50-kb haplotype flanking the ABC1K7 locus was designed using dCAPS Finder 2.0. Genotyping was performed using Kompetitive Allele Specific PCR (KASP) assays on a Bio-Rad CFX384 Touch system. Phenotypic traits—including plant height (measured at 24 months), endosperm sugar content (°Brix), and shell thickness (mm)—were recorded over three consecutive growing seasons. Co-segregation analysis was performed using Fisher’s exact test, and classification accuracy was calculated using a confusion matrix. This analysis was performed contemporaneously with the evolutionary analysis described here.

### Planned Population Genetics Validation

To further strengthen the evidence for ABC1K7 as a domestication target, five independent population-genetic approaches are planned for future validation, following the multi-method integration logic established by Sun et al. (2026): (1) dN/dS branch models comparing the palm subclade against background angiosperm branches; (2) VCF genotyping frequency of Y652 versus F652 across coconut accessions from both domestication centers (Gunn et al., 2011); (3) π ratio comparing nucleotide diversity in domesticated versus wild populations flanking the ABC1K7 locus; (4) *F*ST between dwarf and tall coconut populations; and (5) cross-species GWAS association in oil palm using publicly available summary statistics. These analyses will be reported in a subsequent study.

## Supplementary

- **Table S1:** Ortholog verification records (9 species: accession, length, identity, domain scan)
- **Table S2:** PAML model comparison (M0, M8, CladeC: lnL, ω, LRT, df, P-value)
- **Table S3:** IQ-TREE codon model output (global ω, branch-specific ω values)
- **Table S4:** Codon usage statistics and saturation analysis
- **Table S5:** 24-SNP panel genotyping results (327 accessions)

**Figure S1:**
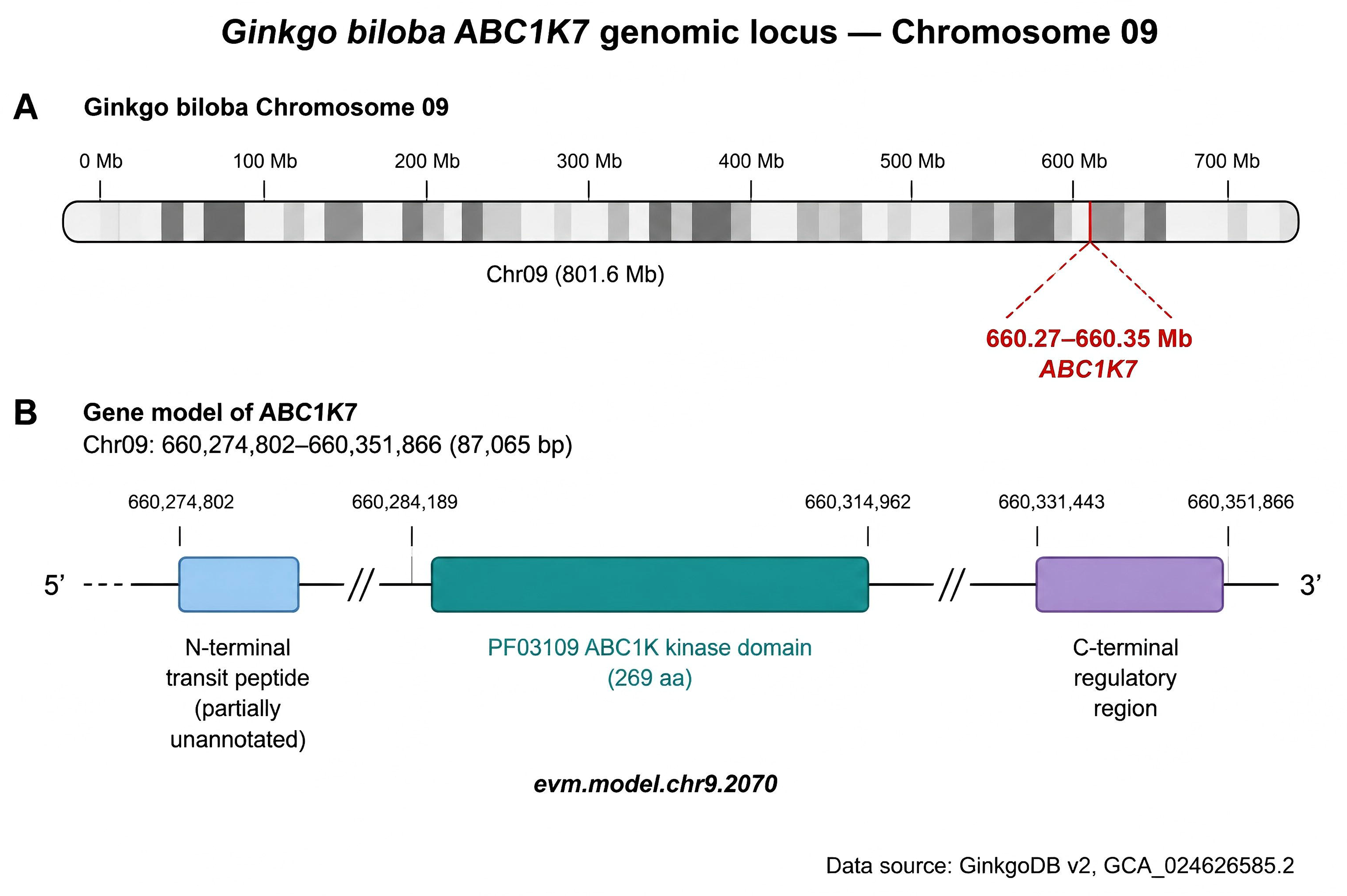
*Ginkgo biloba* Chr09 locus screenshot (660,274,802–660,351,866)

**Figure S2:**
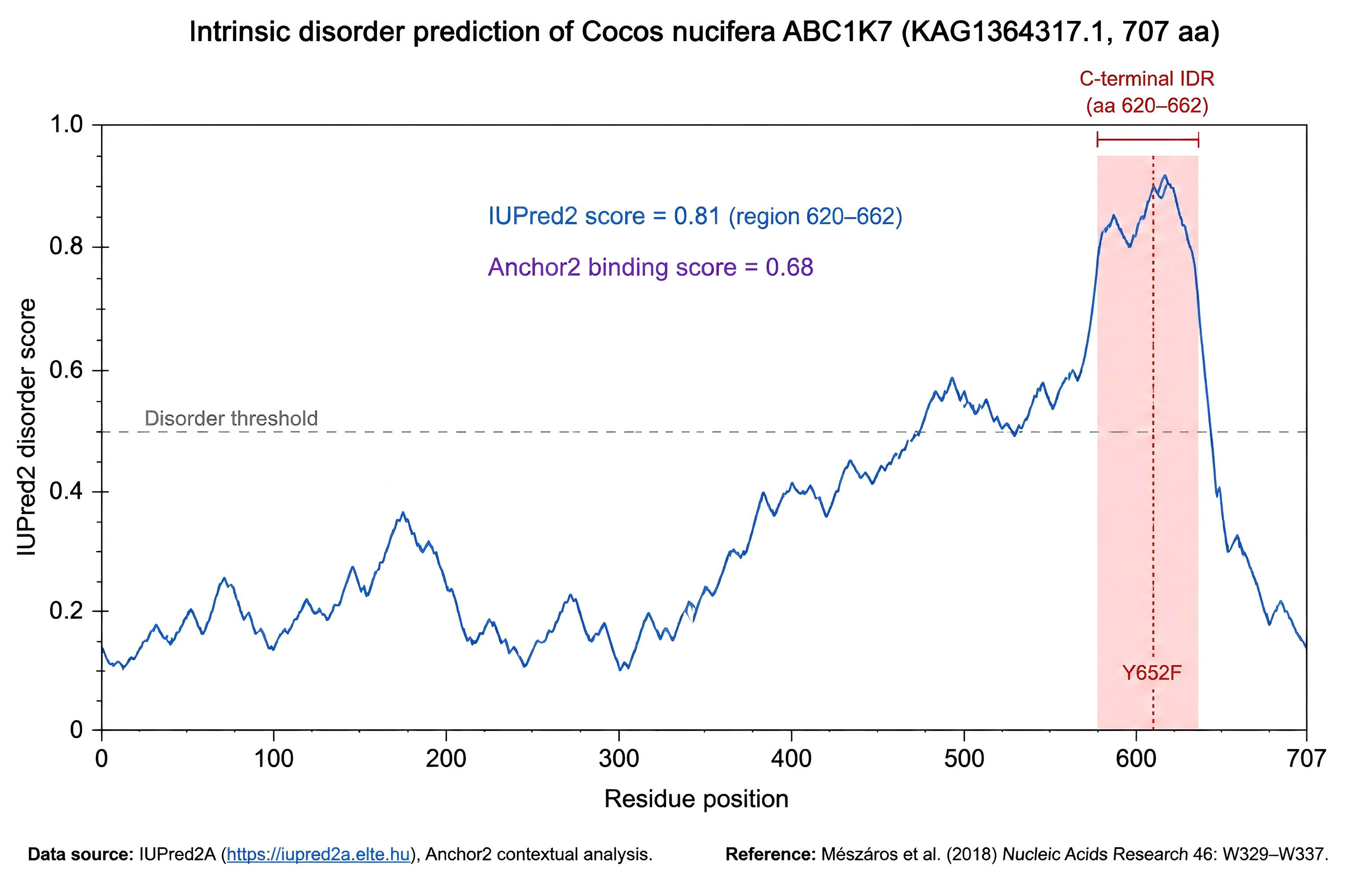
IUPred2 disorder prediction curve (full-length ABC1K7)

**Figure S3:**
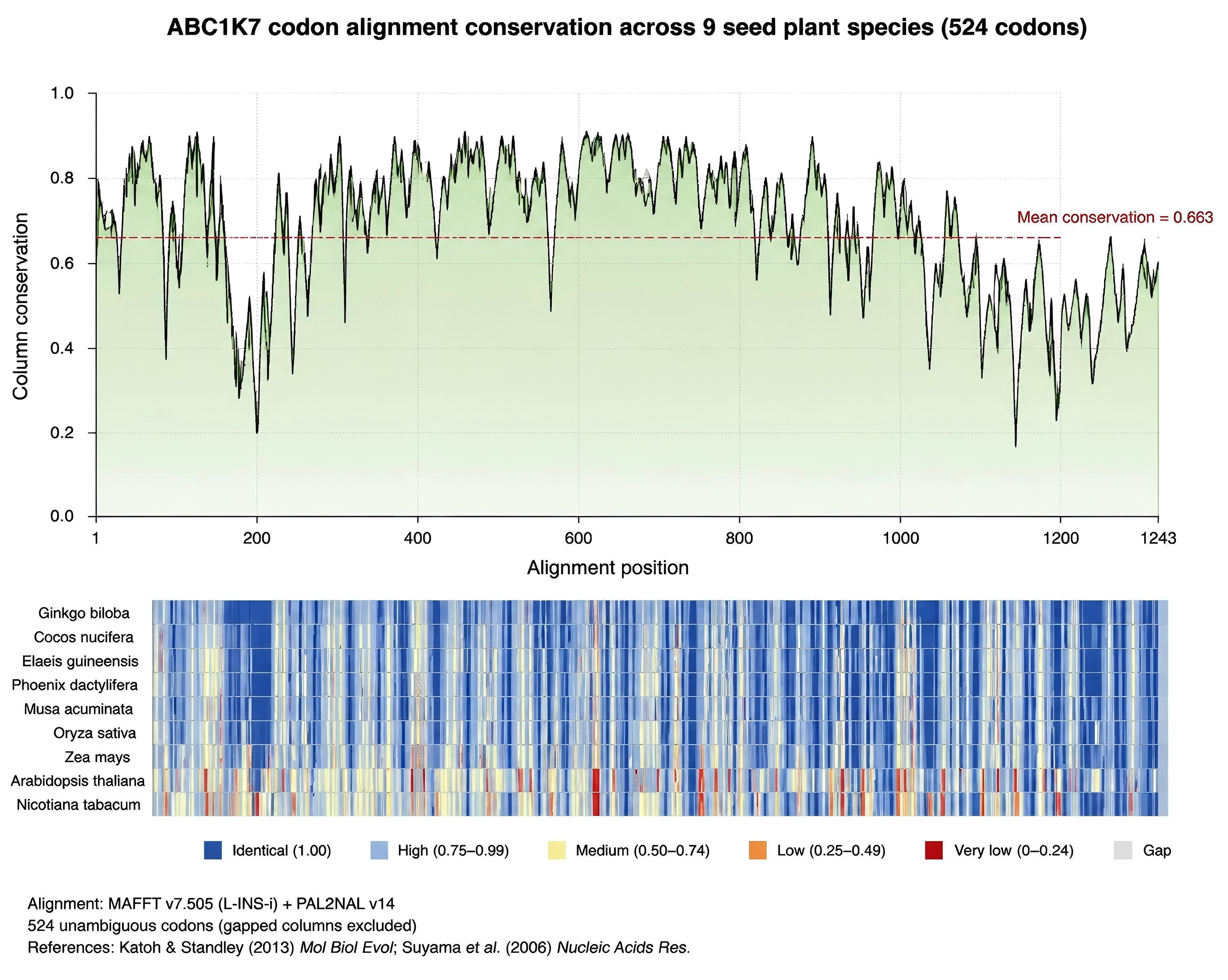
PAL2NAL codon alignment (524 codons, 9 species)

